# An Investigation of the Mechanism for a Novel Influenza Inhibitor Designed to Target the Import of PB1

**DOI:** 10.1101/2021.03.05.434155

**Authors:** Alexander Dorius, David D. Busath

## Abstract

A novel inhibitor of PB1 import, referred to here as GM30, previously displayed nuclear retention of viral nucleoprotein (NP), which spurred the question of whether the compound directly or indirectly blocked viral ribonucleoprotein (vRNP) complex export. vRNPs are exported from the nucleus via exportin 1 (XPO1). Using verdinexor (VNXR), a direct inhibitor of vRNP export that binds to XPO1 and also blocks nuclear factor kappa B (NF-κB) export, we found that GM30 does not block nuclear export of NF-κB. GM30 likewise demonstrated high nuclear retention of NP but not as much as the direct inhibition of nuclear export by VNXR. When the compound was added hours after infection, the compound lost its ability to block nuclear export but VNXR retained its ability to block nuclear export. GM30 is therefore likely an indirect inhibitor of nuclear export because it disrupts the vRNP complex formation and impedes export from the nucleus by reducing nuclear import of PB1.

## 1. Introduction

The influenza virus kills ∼400,000 people around the world each year [1]. Influenza’s high rate of mutagenicity renders vaccination only a partial prevention of influenza infection [2]. Alternative, molecule-based treatments for influenza have focused on targeting viral factors; i.e., M2 channel inhibitors, neuraminidase inhibitors, RNA dependent RNA polymerase complex inhibitors, etc. [3-5]. However, these inhibitors are often vulnerable to development of resistant mutations [6]. But, novel inhibitors that target highly conserved vulnerabilities within the virus’ proteome may provide opportunities for resistance-invulnerable molecule-based therapies [7]. Also, novel therapeutics that target viral-host interactions [8] are especially promising because any viral mutations that might prevent drug binding at the interface between a virus protein and a host protein is likely to disable the natural host interaction as well. Mohl et. al sought out a novel method of inhibition, similar to the ivermectin interference with importin chaperoning of vRNPs into the nucleus [9], but targeting instead importin chaperoning of newly transcribed viral polymerase basic 1 (PB1) at its host importin Ran Binding Protein 5 (RanBP5) binding site to block the formation of the PB1-polymerase acidic(PA)-RanBP5 complex for nuclear import and vRNP completion [10].

They found that compound **20** (referred to here as GM30), derived from a compound identified by library docking at a likely PB1 binding site, is a potential lead compound [10]. GM30 demonstrated low micromolar efficacy, a selectivity index of 6.4, and no resistance formation in passaging experiments. [10]. Its slightly smaller parent compound showed a good fit in simulations of docking to PB1 from both influenza A and B with good positioning to inhibit interactions with RanBP5 by a foot-in-the-door mechanism. During the localization assays performed by Mohl et al., GM30 successfully blocked nuclear import of PB1; however, it also stimulated retention of viral nucleoprotein (NP) in the nucleus [10]. These results left an open question as to how GM30 was blocking the export of NP. Mohl et. al attributed the reduced nuclear export of NP to an indirect effect, brought about by the reduced presence of PB1 needed for vRNP completion in the nucleus. This is plausible because viral polymerase, assembled in the infected cell nucleus, transcribes its own vRNA strand, which aggregates with viral NP as it forms, to form vRNPs, which are then exported [11-14]. Hence, without PB1, NP could accumulate in the nucleus. Other molecules such as Leptomycin-B (LMB) and VNXR [15, 16], demonstrate similar nuclear retention of NP, but through a direct mechanism, i.e. by binding to exportin 1 (XPO1), which shuttles the vRNP complex across the nucleus and into the cytoplasm for subsequent virion progeny [17]. In fact, GM30 bears a superficial resemblance to VNXR (**Figure 1**). Perhaps GM30 directly blocks vRNP export in addition to its inhibition of PB1 nuclear import.

**Figure 1.**
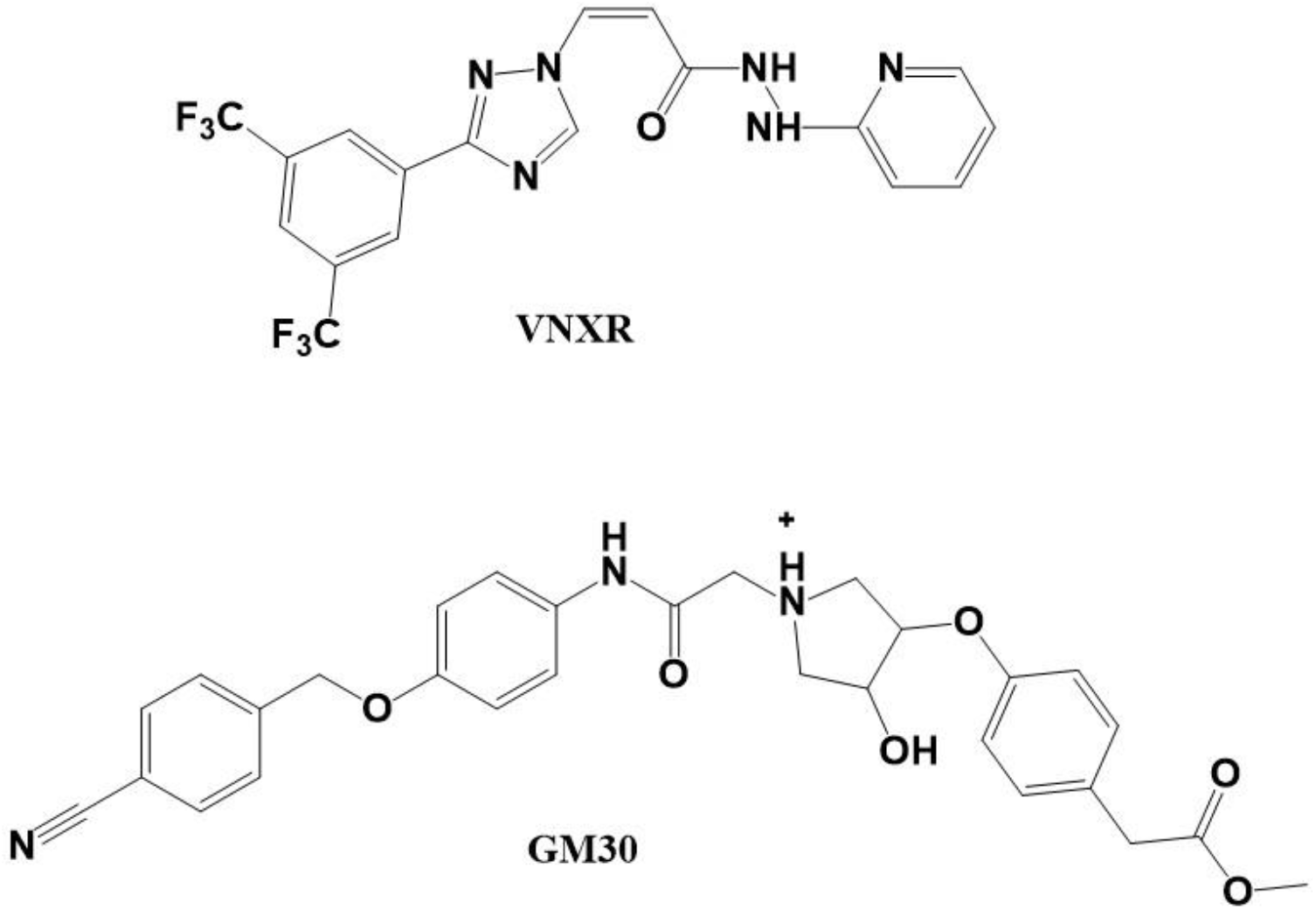
Verdinexor (VNXR) and GM30 share superficial resemblances in structure, positioning of high electron density groups, and possible pi-bonding interaction sites.

Here we report that GM30 does not affect the nuclear export of NF-κB. The timing and impact of GM30 on nuclear localization of NP are contrasted with those of VNXR. Results indicate that GM30 inhibition of PB1 import indirectly inhibits the nuclear export of NP, as previously surmised, by inhibiting the complexation of NP into vRNPs, which depend on nuclear PB1 to serve as a nidus for their formation.

## 2. Materials and Methods

### 2.1 Cell Cultures and Influenza Virus

Madin-Darby canine kidney (MDCK) cells were cultured in Dulbecco’s modified Eagle’s medium (DMEM) supplemented with 5% fetal bovine serum (FBS) and 100 mg/ml of streptomycin. Cells were incubated at 37°C with 5% CO_2_. The Influenza virus strain A/CA/2009 (H1N1) was harvested from inoculated MDCK cells after 48h incubation, and the multiplicity of infection (MOI) was determined using FITC-anti-influenza A polyclonal antibody (Millipore, Billerica, MA) with fluorescence microscopy and cell-counting software after inoculating MDCK cells at varying titers for 18h.

### 2.2 NF-κB Localization Immunofluorescence

MDCK cells were seeded onto 12mm glass coverslips; treated with 3 µM VNXR (Cayman Chemical Company, Ann Arbor, MI) or 30 µM GM30 (a gift from Dr. David Michaelis, BYU) in 1% DMSO, or with vehicle (1% DMSO) and incubated at 37°C for 4h [16]. Coverslips were washed, fixed for 15min with 4% paraformaldehyde in phosphate buffer solution (PBS), and stained with rabbit monoclonal antibody to the p65 subunit of NF-κB (Cell Signal Technology, Beverly, MA) antibody overnight at 4°C. After, the sample was incubated with Alexa 546-conjugated goat anti-rabbit (Invitrogen, Carlsbad, CA) secondary antibody at RT for 2h and stained with 4’,6-diamidino-2-phenylindole (DAPI) before mounting onto slides with Fluoromount-G (Southern Biotech, Birmingham, AL).

### 2.3 NF-κB Localization Imaging and Analysis

Samples were imaged using an Olympus Fluoview FV1000 confocal microscope at 10X magnification, 1600 by 1600 pixels, and 20µs/pixel. An Argon 405nm laser diode excited the DAPI, and a 543nm Helium-Neon laser excited the Alexa 546-conjugate. CellProfiler was used to analyze images for their respective cell count, Pearson correlation coefficients (PCC), and merge channels [18]. For cell count, nuclei were counted using the “IdentifyPrimaryObjects” module with nuclei measuring 20-70 pixel units, global thresholding, minimum cross entropy, clumping allowed, and dividing lines allowed. For PCC, colocalization of NF-κB and nuclei was measured using the “MeasureColocalization” module at a 15% threshold of maximum intensity and across the entire image. Adjusting the threshold to 7% made no change in PCC value, signifying that extracellular noise was mitigated and did not produce an over-inflated PCC value despite using the entire image [19]. At least 6 images per treatment were taken, and the average PCC value and respective standard error of the mean (SEM) was calculated. Statistical significance was calculated and compared using the student’s t-test based on the linear relationship between probe localization and the fluorescence measurement [19].

### 2.4 Nucleoprotein Localization Assay

MDCK cells were seeded onto glass coverslips, washed with DMEM without FBS, and soaked in 10 µM VNXR-, 50 µM GM30-, or vehicle-treated media for 2h before inoculation. Viral media was harvested from MDCK cells infected at 0.1 MOI for 48h and then treated with tosyl phenylalanyl chloromethyl ketone (TPCK) treated trypsin at 1 µg/mL (Sigma) for 30min at RT. After drug pre-soak, drug-treated media was decanted, and chilled viral- and drug-treated media was added and then incubated at 4°C for 1h to maximize synchronization of infected cells [16, 20]. Viral media was then decanted, cells were washed with cold media, and then incubated with drug-treated media at 33°C to initiate cell infections. After time points 12hpi and 16hpi, the cells were washed with PBS, fixed with chilled 4% Paraformaldehyde in PBS for 15min, dried, and then stained for 45min at 37°C in a humidified chamber with both DAPI and with FITC-anti-influenza A polyclonal antibody (Millipore, Billerica, MA) primarily effective for binding viral NP. Non-adherent antibody and DAPI were washed off with PBS containing 0.05% Tween20, 0.5 mM MgCl and 3.1 mM sodium azide and distilled water. Coverslips were then mounted on slides using Fluoromount-G (Southern Biotech, Birmingham, AL).

### 2.5 Nucleoprotein Localization Visualization and Analysis

Cells were scored using an Olympus Fluoview FV1000 confocal microscope at 20X magnification, 1600 by 1600 pixels, and setting the acquisition control to X4 Focus. Cells with low intensity for NP were excluded to focus on synchronized infections. The Z-plane of the image slice was adjusted until maximum NP intensity was achieved relative to the whole region of interest. Cells scored as nuclear (N) had >75% NP localized to the nucleus. Cells scored as cytoplasmic (C) had >75% NP localized to the cytoplasm. All other cells were scored as nucleocytoplasmic (NC) [21]. To maximize the power of the study, samples were scored blindly, infected cells were selected at random as they came into microscopic view, and at least 80 cells per sample were scored. Several samples were rescored to confirm rater consistency.

### 2.6 Nucleoprotein Localization Late-Intervention Assay

The nucleoprotein localization method was used in a subsequent experiment, with the only change made being the removal of the pre-soak with each respective drug and instead the samples being treated with drug starting at 4hpi or 7hpi and continuing until fixation/visualization at 12hpi. This late-intervention paradigm was designed to explore whether GM30 will retain viral NP in the nucleus after hypothetical import of PB1 has already occurred. In theory, if GM30 blocks PB1 import directly, but does not also block NP export directly, it should not impact NP nuclear localization in these late-intervention studies because PB1 activated vRNP formation should occur on schedule. In contrast, VNXR is expected to block the vRNP export in these late-intervention studies because vRNP export occurs later in the infection cycle than PB1 import [20]. We used two intervention delays to help identify the delay time for optimal contrast.

## 3. Results and Discussion

### 3.1 NF-κB Localization Demonstrating Possible Interactions with XPO1/CRM1

VNXR is known to interact with the host protein XPO1, also known as chromosomal maintenance protein (CRM1), inhibiting the export of vRNPs, which utilize XPO1 for nuclear export [16]. Selinexor and VNXR have also demonstrated the capability to inhibit the nuclear export of NF-κB, which also utilizes XPO1 [22, 23]. As demonstrated by Mohl et. al, cells treated with GM30, expected to block PB1 import by binding to RANBP5, also exhibited retention of viral NP in the nucleus; an indirect mechanism for inhibiting the export of vRNPs was hypothesized, namely mass action (lack of PB1 leads to low production of exportable vRNPs and thus accumulation of viral NP in the nucleus) [10]. Perhaps instead, or in addition, GM30 also inhibits the protein-protein interaction with XPO1 and the vRNP cargo complex similarly to VNXR [17]. If true, GM30 would also be expected to inhibit other similar XPO1 functions such as the export of NF-κB from the nucleus.

Localization of NF-κB assays were styled after those performed by Perwitasari et. al, using GM30 as the treatment of interest, VNXR as the positive control, and vehicle (1% DMSO) as the negative control [16]. By qualitative assessment, VNXR demonstrated nuclear retention of NF-κB as seen in **Figure 2B**, and GM30 showed no distinguishable results when compared to vehicle. As seen in **Figure 2A**, VNXR had a PCC, or coefficient of NF-κB colocalization with nuclear stain (DAPI), of 0.69 ± 0.02, which demonstrated high overlap between NF-κB and the nuclei. GM30 had a low PCC of 0.27 ± 0.02 not different from vehicle (0.27 ± 0.03). VNXR significantly differed (P<0.00001) from GM30 and DMSO. Based on these results, GM30 does not interact with XPO1 to inhibit the nuclear export of NF-κB, whereas VNXR does.

**Figure 2.**
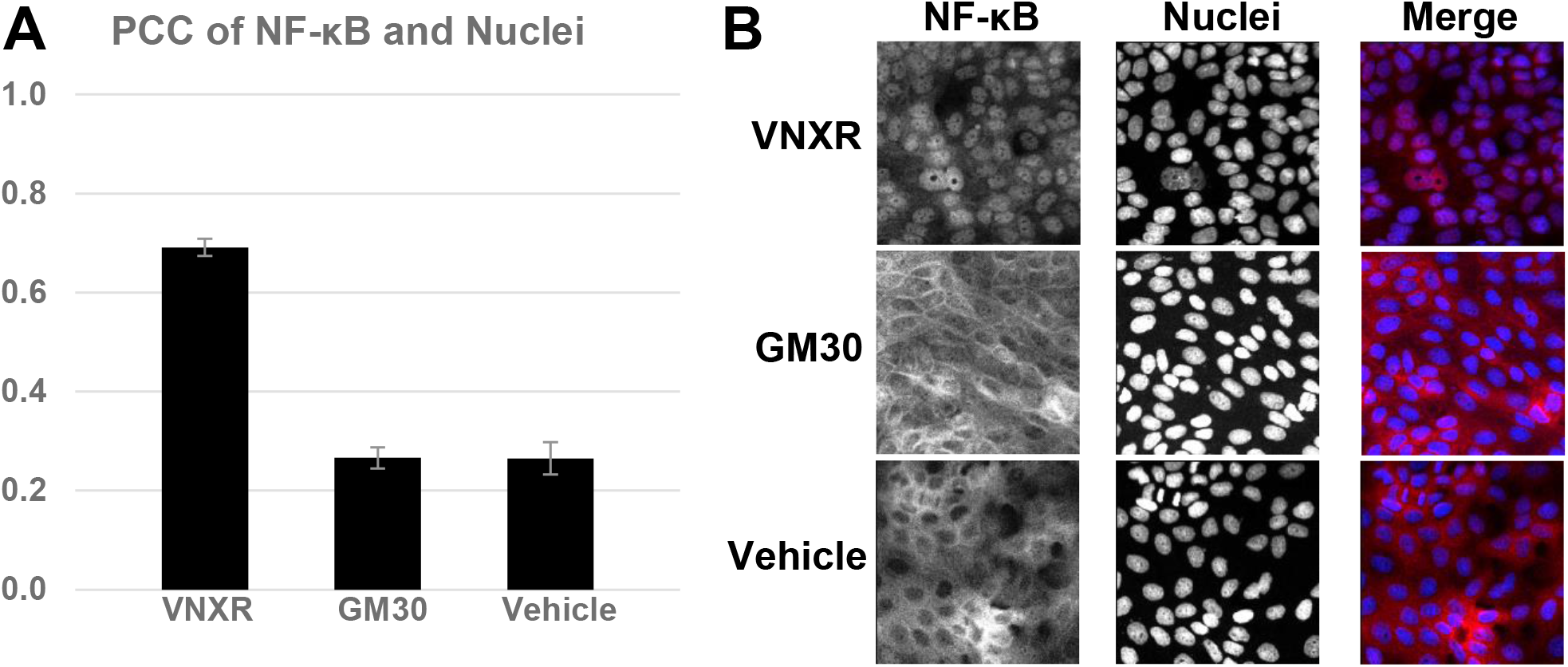
(**A**) Pearson colocalization coefficients (PCC) given for MDCK cells treated with 3 µM VNXR, 30 µM GM30, and 1% DMSO for 4 h. Cells were fixed, stained for the NF-κB p65 subunit and for the nuclei, and imaged using confocal fluorescence microscopy. Images containing ∼10^3^ cells per image were analyzed with the colocalization module CellProfiler to generate PCCs. (**B**) Images of each treatment group with associated channels containing Alexa-546 (NF-κB) and DAPI-stained nuclei (Nuclei) were merged using CellProfiler (Merge). NF-κB probe is labeled red, and Nuclei are labeled blue in Merge.

### 3.2 Viral Nucleoprotein Localization

Mohl et. al conducted viral NP localization experiments in tandem with PB1 localization; however, the infected cells were heterogeneous in their localization, with some showing high nuclear localization of NP, some high cytoplasmic localization, and some with both nuclear and cytoplasmic NP at each time point examined [10]. To achieve more homogenous results, the cold inoculation method was utilized to improve synchronization of the virus infection cycles [16, 20]. Other NP localization experiments have noted difficulties in achieving homogenous results [20, 24], and we, too, still had considerable heterogeneity, though there seemed to be some improvement. Also, we hoped the reproduction of Mohl’s NP localization experiment would allow comparison with VNXR using quantitative analysis. This comparison could help distinguish whether GM30 localizes viral NP to the nucleus directly or indirectly.

GM30 produced substantial NP nuclear localization, although not as great as VNXR for both 12hpi and 16hpi time points (**Figure 3**). GM30-induced nuclear localization (0.42 ± 0.05) was significantly higher (P<0.001) than vehicle (0.20 ± 0.04) 12hpi, and substantially higher (0.71 ± 0.05 vs. 0.08 ± 0.03, P<0.00001) at 16hpi. This increase was not as great as that produced by VNXR, however, at either time point: 0.69 ± 0.03 at 12hpi and 0.94 ± 0.02 at 16 hpi.

**Figure 3.**
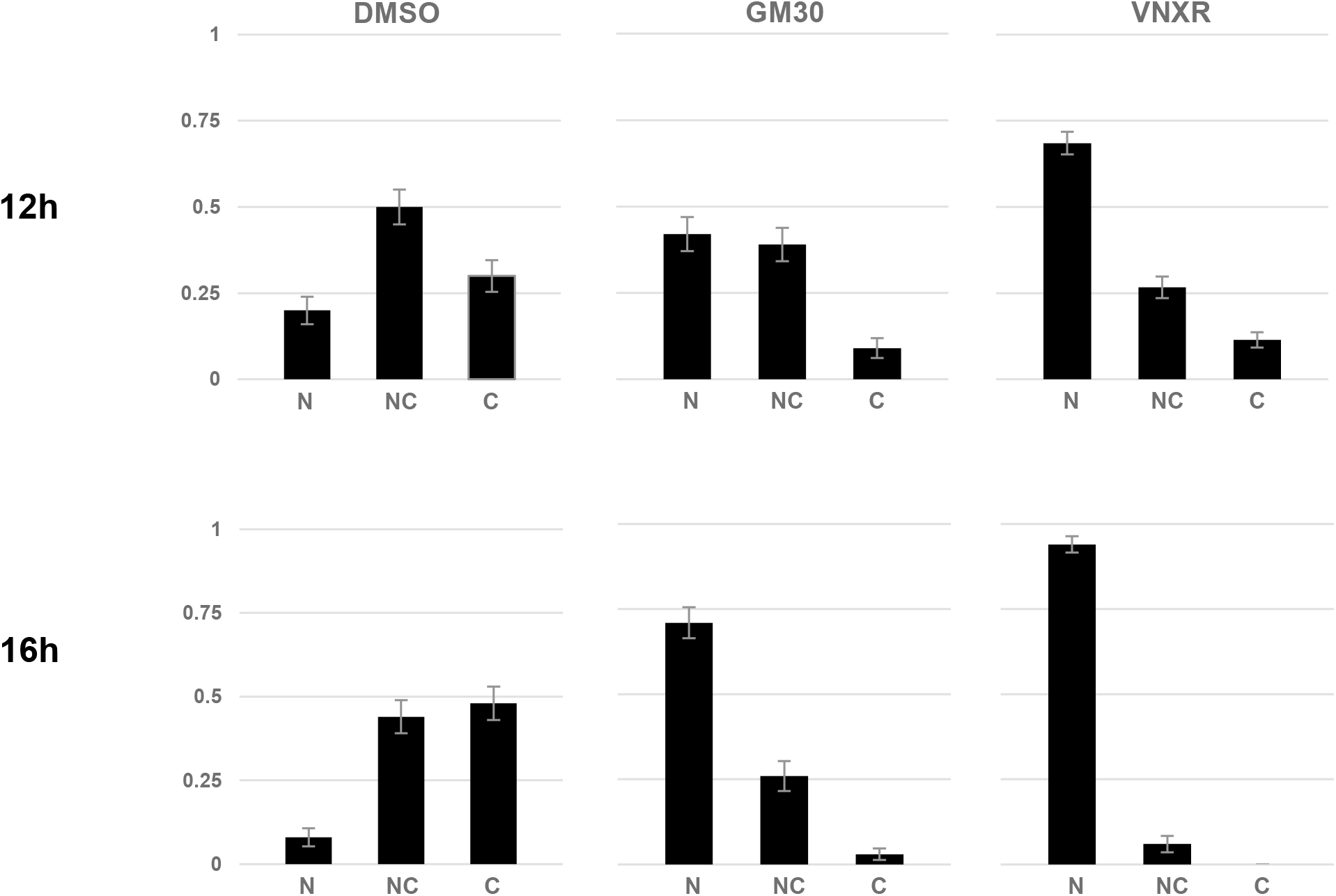
MDCK cells were pre-treated with 10 µM VNXR, 50 µM GM30, and 1% DMSO for 2h before inoculation. At inoculation, the cells were infected with virus at an MOI of 0.1 in chilled, drug-treated media and incubated at 4°C for 1h. Viral media was decanted, cells washed with chilled media, and then incubated at 33°C with drug-treated media until their respective endpoints. At 12h and 16h post infection, cells were fixed, stained for viral NP and nuclei, and scored blindly for localization of viral NP using confocal fluorescence microscopy. Localization was scored qualitatively as >75% nuclear (N), >75% cytoplasmic (C), or mixed (NC) for at least 80 cells/sample.

Interestingly, vehicle exhibited an increase in the fraction of cells with primarily cytoplasmic (C) viral NP localization (compared to the two drugs) at 12 hpi (0.30 ± 0.05) and more so at 16hpi (0.48 ± 0.05). This is to be expected in a cell in the late stage of infection where the vRNPs, containing most of the NP, have been exported into the cytoplasm for packaging and viral exosis [25, 26]. This did not occur for GM30 or VNXR, which both show very low primarily cytoplasmic distribution, especially at 16 hpi. The decrease in cytoplasmic localization at 16 hpi for the two drugs, with concomitant increased fraction of nuclear localization, gives the impression that cells with strong nuclear localization may persist in culture, while cells with failure to maintain nuclear localization proceed to export virus and involute by 12hpi with increasing involution by 16hpi. Similar trends with VNXR were exhibited when qualitative assessment of NP localization was performed by Perwitasari et. al [16].

The significantly greater nuclear localization of NP produced by VNXR compared to GM30 at 12 hpi (P<0.00001) and 16 hpi (P<0.00001) likely manifests differing mechanisms between VNXR and GM30. In addition, the analyses performed by Mohl *et al*. indicate a difference in mechanisms [10]. They proposed that nuclear retention of viral NP may occur because vRNP formation in the nucleus relies on the nuclear presence of PB1, which is blocked for nuclear import by GM30 [10]. Without nuclear import of newly synthesized PB1, there would be no seeding or completion of NP- and RNA-containing vRNP complexes by complete polymerase trimers [27]. Our data seem to rule out the alternative hypothesis that GM30 may interact with XPO1 to inhibit vRNP export because GM30 is much less effective at causing nuclear localization of NP than is VNXR, a known XPO1 blocker. Combined with our finding above that GM30 fails to block NF-κB export via XPO1 while VNXR is effective, it appears that nuclear localization of NP is likely due to an indirect inhibition effect rather than the direct inhibition of vRNP nuclear export.

### 3.3 Viral Nucleoprotein Localization with Post-Infection Drug Treatment

To better distinguish the mechanism of GM30 from that of VXNR, a time-of-exposure study was designed in which each drug was added later in the viral cycle, either beginning at 4hpi where PB1 import is expected to have already begun or 7hpi where vRNP export is beginning to start [16, 20]. Viral globular proteins translated from nascent viral mRNA appear to begin synthesis and nuclear import starting as early as 4hpi [20]. Also, vRNP nuclear export appears to begin around 8hpi and peaks around 12hpi [16, 20, 27]. According to Mohl’s indirect-effect hypothesis, treatment with GM30 at 4hpi should allow a short influx of PB1 proteins to enter the nucleus, reducing the ability of GM30 to induce the indirect nuclear localization of NP. Whereas GM30 treatment at 7hpi should allow significant PB1 import so that it could seed production and nuclear export of vRNPs, substantially reducing or eliminating the indirect nuclear localization of NP. Although drug is added at 4hpi or 7hpi, there is likely a delay in action based on the time needed for the drug to permeate the cell membrane. In contrast, assuming it is reasonably permeant, VXNR would be expected to block the later event of vRNP-export from the nucleus at both exposure starting-points, producing nuclear localization of NP in the form of vRNP complexes. Therefore, assuming the epitope for the polyclonal anti-influenza A antibodies we used are exposed in both globular and vRNP forms of NP, we tested whether nuclear localization of NP was higher in the 4hpi GM30 exposure study than in 7hpi GM30 exposure study.

The indirect-effect hypothesis was supported by the results of this late-treatment experiment as well. Application of GM30 (**Figure 4)** at 4hpi (0.16 ± 0.04) or 7hpi (0.25 ± 0.04) eliminated (P<0.05) the increased NP nuclear localization seen with pre-soaked cells (0.42 ± 0.05, **Figure 3**) relative to vehicle. VNXR applications starting at 4hpi (**Figure 4A**) and 7 hpi (**Figure 4B**) were not significantly different from the original (**Figure 3**) as expected. VNXR continued to exhibit strong inhibition of nuclear export with proportions at 0.67 ± 0.05 and 0.63 ± 0.05, at 4hpi and 7hpi, respectively. It was surprising that nuclear localization was higher after GM30 treatment at 7 hpi than after treatment at 4 hpi, when under the indirect-effect hypothesis it was expected to be lower. We attribute this to the rising propensity of nuclear localization with time in an unimpeded infection, as demonstrated by the results with the vehicle, suggesting that PB1 entry beyond 7hpi is still important for final vRNP formation in and clearance from the nucleus.

**Figure 4.**
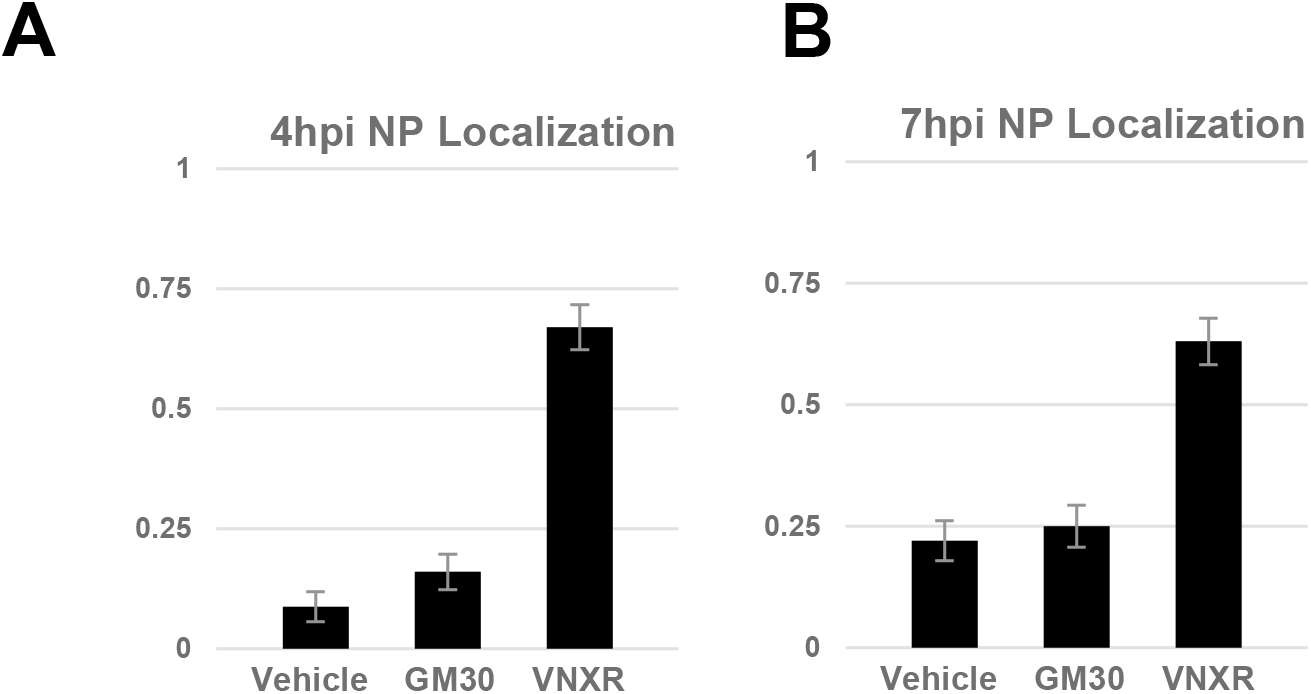
MDCK cells were infected with virus at an MOI of 0.1 in chilled, drug-treated media and incubated at 4°C for 1h. Viral media was decanted, cells washed with chilled media, and then incubated at 33°C until intervention treatment. Media was replaced by 10 µM VNXR-, 50 µM GM30-, or vehicle (1% DMSO)-treated media at intervention time points **A**) 4hpi (4hpi NP Localization) or **B**) 7hpi (7hpi NP Localization). At 12hpi, cells were fixed, stained for viral NP and nuclei, and scored blindly using confocal fluorescence microscopy. Nuclear localization of viral NP (NP Localization) was scored for at least 80 cells per treatment and compared.

These results confirm and further elucidate GM30’s probable indirect-effect mechanism. Because VNXR remained effective when administered late while GM30 was effective only when administered early, the direct effect of GM30 on vRNP by XPO1 as a cause of nuclear localization of NP was successfully ruled out. This leads to the conclusion that GM30 produces nuclear NP localization by the indirect-effect mechanism and failure to seed and complete vRNPs with the polymerase complex required for nuclear NP export [28]. From its failure to block NF-κB export, GM30 does not directly appear to interact with the nuclear export pathway characterized by VNXR’s interaction with XPO1. Rather GM30 likely stops the assembly of vRNPs through its ability to block PB1’s entry into the nucleus.

In addition, our results further characterize the need of the trimeric polymerase complex to properly assemble vRNPs in the nucleus. Replication of influenza, negative-sense viral ribonucleic acid (vRNA) requires a positive-sense template, referred to as complementary RNA (cRNA)-containing RNP complexes (cRNP) [29]. The cRNP is used as the template to create progeny vRNPs in the nucleus after nuclear import of the necessary nascent viral proteins, NP, PA, PB1, and PB2 [29]. However, for replication to proceed, the nascent vRNP must form its trimeric polymerase complex before beginning *trans* replication of the cRNA on the cRNP [11-13]. As *trans* replication via the nascent trimeric polymerase complex of the assembling vRNP proceeds, NP is recruited to the growing segment of replicated vRNA protecting the vRNA from degradation [14]. After complete assembly of the vRNP, the vRNPs are then exported out of the nucleus with the aid of XPO1 and many other factors [30, 31]. The indirect-effect hypothesis for how GM30 produces an increased fraction of cells with NP nuclear localization is based on the concept that without the needed formation of the trimeric polymerase complex for vRNP assembly, NP would accumulate in the nucleus without the usual export as a component of vRNPs. Recruitment of NP for vRNP assembly acts as a sink.

If vRNP assembly is perturbed, NP remains in its globular form in the nucleus which likely accumulates over time. Since the influx of globular NP is modulated by importin-α proteins and the efflux is only due to passive diffusion [32], GM30 treatment should theoretically produce some degree of NP “leakage” from the nucleus. This explains why GM30 treatment has significantly higher levels of NP nucleocytoplasmic localization than VNXR (**Figure 3**). Under VNXR treatment we do not observe this same effect because vRNP formation still occurs, allowing for vRNPs to mop up excess NP and minimize NP diffusion back into the cytoplasm, with the vRNPs being confined to the nucleus by the block of XPO1. This likewise contributes further to the explanation of why NP nuclear localization is higher for VNXR than GM30.

Interestingly, VNXR is much more effective at producing a high fraction of cells with predominantly nuclear NP when administered late (**Figure 4**). We attribute this to VNXR’s late-stage mechanism, i.e. block of vRNP export, contrasting to GM30’s block of an earlier stage mechanism, import of newly synthesized PB1 to the nucleus. Live single cell imaging with NP state-specific antibodies and other specific complex-inhibiting pharmacological agents [24, 33, 34] might allow these differences to be more completely interpreted in terms of, mass action control of NP complexation inside the nucleus, and chaperoned NP import and export, which ultimately control the timing of NP concentrations inside and outside the nucleus at different stages of the infected cell cycle.

## 4. Conclusion

Mohl et. al discovered a novel inhibitor, GM30, that targeted the PB1 interaction with importin RanBP5 but still exhibited a puzzling accumulation of viral NP in the nucleus. We hypothesized that retention of viral NP seen with GM30 may be due to an additional mechanism, targeting of XPO1. Since XPO1 also exports NF-κB from the nucleus to the cytoplasm, we performed NF-κB localization experiments with both drugs; we concluded that GM30 does not inhibit nuclear export of NF-κB and therefore does not likely interact with XPO1. NP localization for both 12hpi and 16hpi with GM30 and VNXR showed that VNXR fully retains NP in the nucleus more frequently than GM30. Finally, addition of drug after hypothetical import of PB1 into the nucleus, GM30 lost its ability to retain NP in the nucleus while VNXR maintained nuclear retention of NP, confirming that excess nuclear localization of NP caused by GM30 is the result of reduced formation of trimeric polymerase in the nucleus due to absence of PB1. Likewise, vRNP formation is stunted without the source of trimeric polymerase subunits necessary for proper *trans* replication of cRNA on cRNPs. This leaves NP stranded in the nucleus in its globular form instead of assembling into vRNP complexes. The anti-viral mechanism of GM30 is likely to be inhibition of nascent PB1 into the nucleus as predicted.

## Acknowledgements

Chris Mendoza and Dr. Dario Mizrachi helped with preliminary efforts. The research was supported by funding from Brigham Young University.

